# Absence of 8-HDF and MTHF Antenna Chromophore Binding in *Er*CRY4a Suggests a Possible Flavin-Only Cofactor State: Insights from Biochemical and Computational Analyses

**DOI:** 10.64898/2026.02.17.706321

**Authors:** Alisha Bhanu Pattani Ameerjan, Bahareh Dabirmanesh, Jonathan Hungerland, Takaoki Kasahara, Rabea Bartoelke, Glen Dautaj, Ghazaleh Saberamoli, Jessica Schmidt, Jingjing Xu, Ilia A. Solov’yov, Karl-Wilhelm Koch, Henrik Mouritsen

## Abstract

Cryptochromes and photolyases are blue-light-sensitive flavoproteins that generally bind flavin adenine dinucleotide (FAD) and have distinct functions. Cryptochrome 4a (CRY4a) is a protein expressed in the double-cone photoreceptors of the retina in migratory songbirds like European robin (*Erithacus rubecula*) and is hypothesized as the primary sensor for avian magnetoreception. In addition to FAD, most photolyases and some cryptochromes bind antenna chromophores such as 8-hydroxy-5-deazaflavin (8-HDF) or 5,10-methenyltetrahydrofolate (MTHF) to enhance light absorption. Here, we investigated whether *Erithacus rubecula* Cryptochrome 4a (*Er*CRY4a) also binds 8-HDF and/or MTHF. 8-HDF binding was studied by co-expressing *Er*CRY4a with the *fbIC* gene that encodes for 8-HDF synthase and thus for production of 8-HDF in *E. coli*. As a positive control for 8-HDF binding, we expressed *Xenopus laevis* 6-4 photolyase (*Xl*6-4PL) which is known to bind both FAD and 8-HDF. This experiment resulted in successful binding of 8-HDF to *Xl*6-4PL, but not to *Er*CRY4a. We studied the binding of MTHF using *in vitro* reconstitution followed by UV-Vis spectroscopy and isothermal titration calorimetry (ITC) assays. No interaction was observed between MTHF and *Er*CRY4a. To theoretically understand the binding of potential antenna chromophores to *Er*CRY4a, we performed computational analyses. We found no similarity at the relevant binding sites between the sequences of *Er*CRY4a with proteins shown to bind MTHF or 8-HDF. This suggests that the binding pocket is not conserved. Our study proposes that *Er*CRY4a only harbor one light-sensitive cofactor, which in turn suggests a functional specialization different from most photolyases.

## Introduction

The ability of animals to sense and utilize the Earth’s magnetic field for navigation during migration is known as magnetoreception. This remarkable capability has been observed in various species, including for instance songbirds, fish, sea turtles, mammals, and insects ([1–7]. In night-migratory songbirds, the radical pair mechanism involving cryptochrome 4a (CRY4a) as the primary magnetic sensor is the most prominent magnetoreception hypothesis. The radical pair hypothesis proposes that light-induced electron transfer reactions generate radical pairs that are sensitive to weak magnetic fields [8–10]. CRY4a is a light-sensitive protein expressed in the double cone photoreceptors in the retina of migratory songbirds such as the European Robin, *Erithacus rubecula* [11–13]. This protein has a catalytic light sensing chromophore called flavin adenine dinucleotide (FAD) which is located in the C-terminal α-helical domain of a conserved photolyase homology region (PHR domain) [14,15].

Some members of the cryptochrome/photolyase family of proteins are known to bind additional chromophores, mainly 5,10-methenyltetrahydrofolate (MTHF) and 7,8-dimethyl-8-hydroxy-5-deazaflavin (8-HDF) [16–25]. These additional chromophores have been reported to act as photo-antennas capturing photons and transferring light energy to FAD via the Förster resonance energy transfer (FRET) mechanism[16,17].

8-HDF is a deazaflavin derivative that is infrequently synthesized by most host organisms [26]. The biosynthetic pathways for 8-HDF are largely absent outside a few specific groups of microorganisms, algae and mosses. Vertebrates can acquire 8-HDF solely through dietary intake, particularly by consuming algae or mosses that naturally produce this molecule [26–28]. Co-expression of target proteins with 8-HDF synthase encoding genes (either *fbiC or cofG and cofH*) enables *E. coli*, which lacks an endogenous 8-HDF biosynthetic pathway, to produce the 8-HDF chromophore *in vivo* [26]. Two independent studies were conducted on the green alga *Chlamydomonas reinhardtii* animal-like cryptochromes (*Cra*CRY), which function both as a circadian regulating cryptochrome and as a 6-4 photolyase DNA repair enzyme. These studies showed that the binding of 8-HDF as an antenna chromophore to *Cra*CRY kinetically stabilized the flavin-neutral radical (FADH•) and the reduced FAD (FADH^-^) state [17,18]. Furthermore, in *Xl*6-4PL, 8-HDF captures photons and transfers light energy to the active fully reduced FADH^−^ and enhances the DNA repair activity of the enzyme [19]. 8-HDF has also been reported to act as a significant photosensitizer in the photoreactivation process catalysed by DNA photolyase from *Anacystis nidulans* [20,21].

MTHF is a folate derivative that plays a role in the folate metabolism [29]. It is bound to the N-terminal domain of some cryptochrome and photolyase proteins within a specific binding pocket that is exposed to the solvent[22], thereby facilitating exchange of the antenna chromophore, which can lead to its loss during purification [30,31]. Because of this lability, *in vitro* reconstitution by incubating purified protein with MTHF is often required to assess MTHF binding [16]. MTHF has been reported to serve as a key component necessary to transfer energy to FADH^−^, enabling it to enter an excited state and initiate DNA repair in *E. coli* DNA photolyase[25]. Studies on *Ostreococcus tauri* cryptochrome (*Ot*CPF2) and *Phaeodactylum tricornutum* cryptochrome (*Pt*CryP) have highlighted the versatility of MTHF-FAD interactions in improving light energy utilization and electron transfer [23,24].

Xu et al. reported the successful purification and characterization of CRY4a from the European Robin (*Er*CRY4a), a migratory songbird [15]. To date, no second chromophore has been identified in the recombinantly expressed and purified *Er*CRY4a. However, a comprehensive understanding of the function of CRY4a requires investigation of all potential chromophores and their interactions with the protein. In this study, we investigated the potential presence of a second chromophore in *Er*CRY4a. A combination of biochemical, spectroscopic, and computational approaches was used to evaluate whether 8-HDF and/or MTHF binds to *Er*CRY4a. Our results support a model in which *Er*CRY4a operates with FAD as its sole light-sensitive cofactor, indicating a functional specialization that sets it apart from most photolyases.

## Results

### *Er*CRY4a does not bind 8-HDF

All proteins (*Er*CRY4a and *Xl*6-4PL) were expressed and purified with and without 8-HDF synthase. UV-Vis spectroscopic analyses of the proteins *Er*CRY4a and *Xl*6-4PL (as a control) revealed that FAD was present in both. However, a significant difference was observed when comparing the spectral profiles of *Er*CRY4a and *Xl*6-4PL co-expressed with 8-HDF synthase. *Xl*6-4PL co-expressed with 8-HDF synthase exhibited a distinctive spectral signature, characterized by a peak at 440nm and a shoulder at 420nm (Figure 1), indicative of 8-HDF binding together with FAD. *Xl*6-4PL is a well characterized protein that binds to both 8-HDF and FAD and shares high sequence similarity with *Er*CRY4a[19]. Despite this, *Er*CRY4a co-expressed with 8-HDF synthase did not exhibit the characteristic absorption peak of 8-HDF at 440nm (Fig 1D).

**Figure 1:**
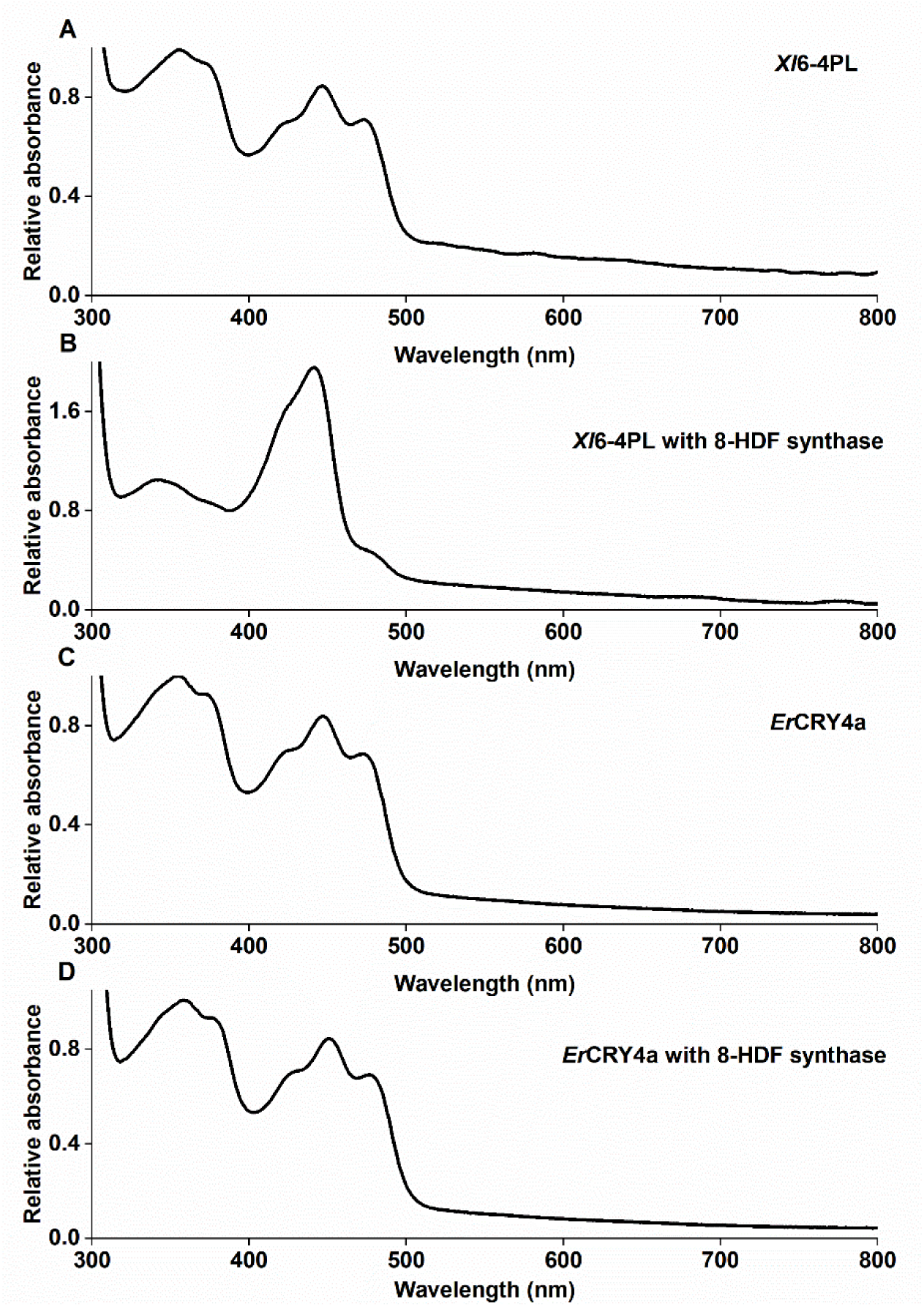
UV-Vis Spectra of *Er*CRY4a wildtype and *Xl*6-4PL expressed and purified with and without 8-HDF synthase. Spectra were recorded using 30 μM *Er*CRY4a (for both with and without 8-HDF synthase samples) and 16 μM *Xl*6-4PL (for both with and without 8-HDF synthase samples). Relative absorbance against wavelength(nm) was plotted by normalizing each spectrum to the absorbance at 355nm. A) *Xl*6-4PL expressed without 8-HDF synthase shows the characteristic FAD absorption spectrum B) *Xl*6-4PL co-expressed with 8-HDF synthase displays the expected 8-HDF chromophore signature, with a prominent peak at ∼440 nm and a shoulder near 420 nm, in addition to FAD C) *Er*CRY4a expressed without 8-HDF synthase exhibits the typical FAD absorption profile D) *Er*CRY4a co-expressed with 8-HDF synthase shows no additional absorption features corresponding to 8-HDF, indicating that *Er*CRY4a does not bind 8-HDF under these expression conditions.

### *Er*CRY4a shows no detectable interaction with MTHF

To investigate the potential interaction between MTHF and *Er*CRY4a, *in vitro* reconstitution followed by UV-Vis spectroscopy and isothermal titration calorimetry (ITC) was performed. 16μM *Er*CRY4a and 800µM MTHF were incubated for one hour in the dark at 4°C. UV-Vis spectroscopy analysis of *Er*CRY4a following *in vitro* reconstitution with MTHF revealed a characteristic absorbance of free MTHF at 360 nm (Figure 2) prior to washing using centrifuge filters, indicating that MTHF remains unbound in the mixture. Bound MTHF in cryptochromes/photolyases has a red shift and absorbance around 380nm[16,22,23]. After filtration with 30 kDa centrifuge filters, the free MTHF-specific absorbance was not detectable.

**Figure 2:**
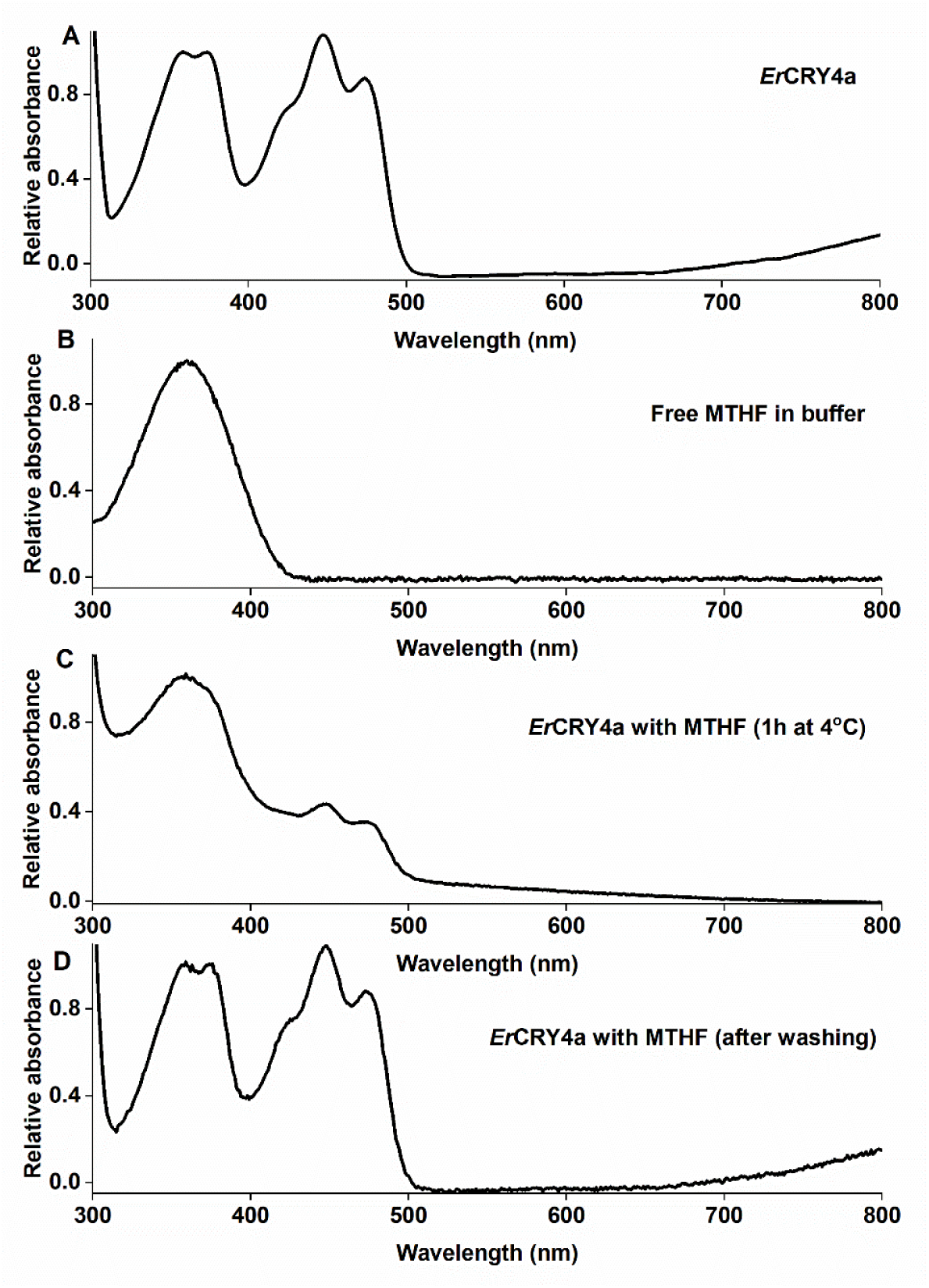
UV-Visible spectroscopy analysis of *Er*CRY4a and MTHF interactions. Spectra were recorded using 16μM *Er*CRY4a, and relative absorbance against wavelength (nm) was plotted after normalizing each spectrum to the absorbance at 360nm. A) *Er*CRY4a alone shows the characteristic FAD absorption peak. B) 800 μM MTHF in buffer exhibits a distinct absorbance of free MTHF at ∼360 nm. C) *Er*CRY4a incubated with 800 μM MTHF for 1 h in the dark at 4 °C shows the FAD peak along with another peak at ∼360 nm, corresponding to free MTHF. D) After washing the *Er*CRY4a-MTHF sample through a 30 kDa centrifugal filter, the 360 nm peak disappears.

### Isothermal titration calorimetry (ITC) confirms a lack of detectable MTHF binding to *Er*CRY4a

To assess the potential interactions between MTHF and *Er*CRY4a, ITC experiments were performed. Prior to the experiment, the instrument underwent initial calibration using a well-established binding pair with 1 mM Ca^2+^ in the syringe and 100 μM EDTA in the calorimetric cell. Titration yielded exothermic heat pulses (µcal/sec) and the corresponding binding isotherm fitted to a one-site binding model demonstrated a distinct saturation point indicative of the binding interactions between the two species (data not shown). Then the interaction of the purified *Er*CRY4a (ranging from 3 to 30 µM) and MTHF (from 400-800µM) at 12 °C and 25 °C was examined. A control titration of MTHF ligand into sodium phosphate buffer at pH 7.5 without *Er*CRY4a was also conducted. Binding of MTHF to *Er*Cry4 would have resulted in either exothermic or endotherm heat pulses approaching the zero baseline at saturation. Notably, the exothermic heat pulses reached similar amplitudes over a wide concentration range and a molar ratio far above a 1:1 stoichiometry. Furthermore, the exothermic heat pulses of the control titration of MTHF titrated to the buffer (left panel) were higher in all the conditions. The resulting integrated heat data displayed in the lower panel of Figure 3A and Figure 3B did not match a typical binding isotherm. Instead, the integrated heat data expressed in kcal/mole of injectant showed scattered profiles and data points are different from zero. The results of the titration might therefore indicate a weak association of MTHF with the protein surface, but not a permanent interaction with the chromophore binding pocket. These findings consistently indicate that there was no detectable interaction between MTHF and *Er*CRY4a under the tested conditions.

**Figure 3:**
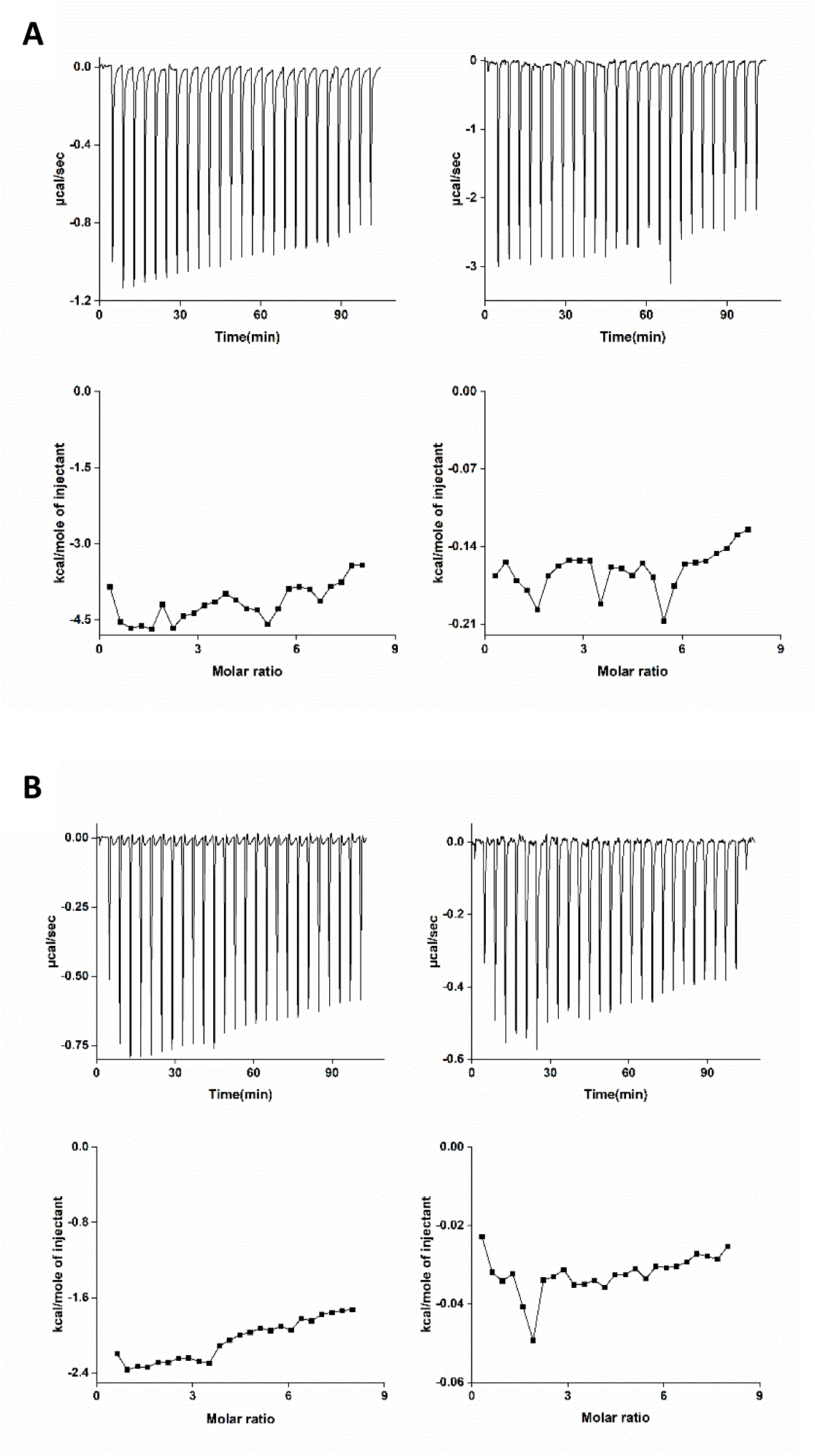
ITC profiles of *Er*CRY4a titrated with MTHF. A) 800µM MTHF was titrated to buffer (left panel) or 16 µM *Er*CRY4a (right panel). Both titrations were performed at 12°C. B) The same titration of MTHF to buffer (left panel) and 16 µM ErCry4a (right panel) done at 25 °C. The data were baseline-adjusted, and the area under the heat pulse curves was analysed resulting in a plot of kcal/mol of the injectant against the molar ratio (lower parts in Figure 3A and B) including the negative controls (MTHF to buffer at both 12 °C and 25 °C) Similar thermograms were observed across a range of *Er*CRY4a concentrations (3–30 µM) and MTHF concentrations (400-800 µM) (refer to Supplementary Figure 1&2).

The combined results from both *in vitro* reconstitution and ITC experiments show that MTHF does not bind to *Er*CRY4a under the conditions tested.

### *Er*CRY4a Shows No Predicted Interaction with 8-HDF and MTHF *in Silico*

As an independent way to assess the potential binding interactions between 8-HDF and MTHF with *Er*CRY4a, the proteins and chromophores we computationally modelled. The fundamental steps of the presented approach are outlined here, while details are provided in the “Methods” section. At the core of the strategy was the positive controls. First, we reproduced that 8-HDF binds to *Xl*6-4PL[19] and that MTHF binds to *Arabidopsis thaliana* cryptochrome 3 *(At*CRY3) [32]. Second, multiple sequence alignments were computed including *Er*CRY4a and all protein structures that were experimentally resolved in the presence of the respective chromophores. The alignment served as an initial probe to estimate where on the protein surface binding could occur and how conserved the amino acid residues were that formed the binding pockets. Afterwards, we employed MD simulations to study the protein conformations in the pH range of 7-8, which yielded a diverse set of realistic conformations. Via molecular docking with VINA, a set of optimized locations for the chromophores were found. During this step, the protein was static, but the protein conformations were accounted for by performing the docking against 3000 different protein conformations sampled from Molecular dynamics (MD). In the docking calculations, the chromophores were allowed to explore the whole protein surface (exhaustive docking) and were not restrained to any predetermined binding pocket. Finally, the docked positions were re-evaluated using additional properties other than the VINA score (a binding ΔG estimate). The diffusion score measures how quickly water molecules would be expected to move, if water molecules were found at the optimized location of the chromophore. The diffusion scores were obtained from the MD simulations in which explicit water molecules were present. The solvent accessible surface area (SASA) score measures the relative change in solvent accessibility when the chromophore is docked into the optimized location. The entropic barrier score estimates in how many directions the chromophore could move out of its optimized location without encountering anything that blocks its path. To average results from similar positions, we clustered the optimized locations such that positions in which the chromophore is in contact with the same amino acid residues are sorted into the same cluster.

The computational results for the analysis of 8-HDF binding are displayed in Figure 4. An excerpt of the multiple sequence alignment in Fig. 4A highlights contacts, primarily but not exclusive formed by hydrogen bonds, between 8-HDF and the surrounding protein structure. The sequence number of *Xl*6-4PL is shown in the top row. The resolved structures *Anacystis nidulans* Photolyase (*An*PL), *Thermus thermophilus* photolyase (*Th*PL), *Chlamydomonas reinhardtii* (*Cra*CRY) and *Synechococcus elongatus* 6-4 photolyase (*Sy*PL) share many similarities in their backbone and sidechain contacts to 8-HDF with the exception that *Th*PL has fewer sidechain contacts. Even though *Xl*6-4PL was not structurally resolved in the presence of 8-HDF and can therefore not be categorized in the same way, the occurrence of R51, D101, E103, R109 and K256 suggests that *Xl*6-4PL exhibits binding patterns that are comparable to those of the resolved structures. Notably, *Er*CRY4a contains charged residues of the same type as most 8-HDF binding proteins at the aligned positions 51, 101, and 103, while lacking those at positions 109 and 256. Therefore, a potential binding motif of 8-HDF in *Er*CRY4a might resemble the one found in *Sy*PL.

**Figure 4:**
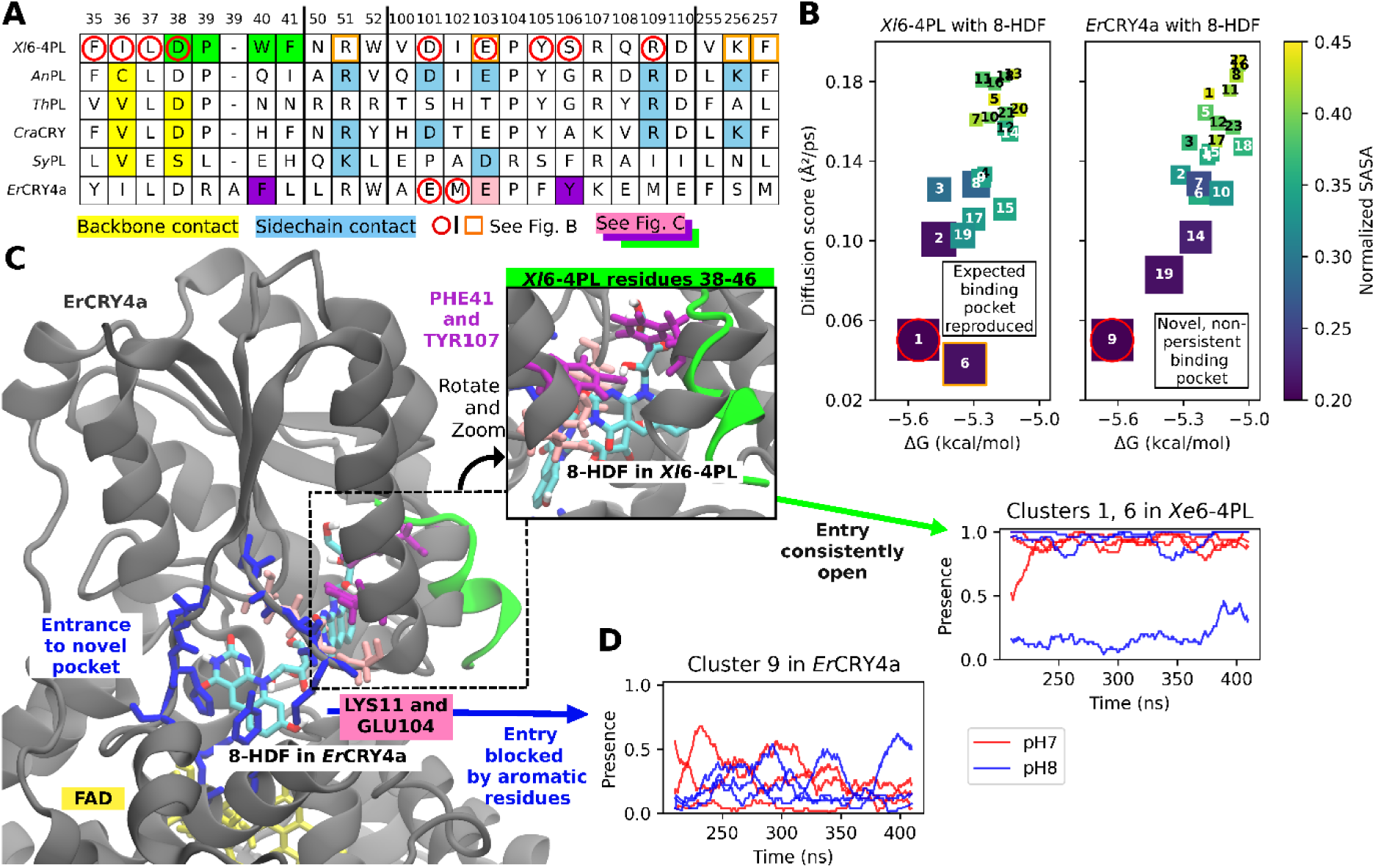
Binding properties of 8-HDF in selected proteins. **A:** Sequence alignments of the crystallized 8-HDF-binders, *Xl6-4PL* and *Er*Cry4a. The numbers in the first row indicate the number of *Xl6-4PL* residues. **B:** Properties of the docking position clusters. The box sizes were scaled with the sixth power of the entropic barrier score. The text color of the cluster numbers is for readability only. **C:** Rendering of the *Er*CRY4a conformation with 8-HDF docked in cluster 9. The 8-HDF position of cluster 1 in *Xl6-4PL* is shown too. The selected residues from *Er*CRY4a are highlighted. The helix that opens the known binding pocket in *Xl6-4PL* is shown in green. **D:** Presence score for the selected clusters over time. Each line corresponds to one MD simulation.

Exhaustive molecular docking calculations involving 8-HDF were performed and the analyzed properties are compiled in Figure 4B. The x- and y-axis show the ΔG score and the diffusion score respectively; clusters are indicated by numbers; box colors show the SASA score and box sizes scale with the entropic barrier. The most likely cluster 1 as well as cluster 6 stand out via the optimal scores across all metrics. As indicated by the red circles in the *Xl*6-4PL row of Fig. 4A, cluster 1 exhibits most contact residues that are expected to occur based on the sequence alignment. K256, however, is not in contact with 8-HDF when the chromophore is docked in the position of cluster 1. By combining the information from sequence alignment, docking and re-scoring, the approach therefore reproduced the expected binding pocket of *Xl*6-4PL.

Performing the same exhaustive docking calculations on the *Er*CRY4a protein conformations revealed viable binding pocket for 8-HDF in cluster 9. All scores of cluster 9 are comparable to those of cluster 1 in *Xl*6-4PL. The corresponding contact residues (see red circles in the *Er*CRY4a row of Figure 4A) are, however, different from those found for the other proteins that contained a bound 8-HDF. Due to unexpected contact residues, the 8-HDF binding pocket of cluster 9 was novel.

Figure 4C shows a rendering of *Er*CRY4a with 8-HDF docked in the position of cluster 9 which can be found in the middle, right behind the amino acids that form the blue-colored entrance to the binding pocket. In the same rendering, another possible position of 8-HDF is shown: the location of 8-HDF in cluster 1 of the *Xl*6-4PL docking results. This hypothetical position is buried behind a grey alpha-helix in the main rendering, and it is more clearly visible in the zoomed-in rendering to the right. Amino acid residues 38-46 of *Xl*6-4PL are displayed in the form of a green helix. Compared to the grey-colored helix of *Er*CRY4a at the same location, the 38-46 residue helix of *Xl*6-4PL is tilted to the right, which opens an entry to the binding pocket. It was calculated how often cluster 1 was among the docked positions in *Xl*6-4PL, which constitutes the presence score shown in Fig. 4D. The entrance to clusters 1 or 6 is continuously open in 5 out of 6 replica simulations of *Xl*6-4PL. The presence score of cluster 9 in *Er*CRY4a, however, was absent in most simulation frames. The partial blockage of the entrance to the pocket of cluster 9 can be attributed to the dynamics of aromatic sidechains within the entrance.

To rationalize the blockage of the expected binding pocket, the protein structures of *Er*CRY4a were combined with the docked positions of 8-HDF produced by docking 8-HDF to *Xl*6-6PL and steric clashes were counted. It was found that the unavailability of the binding pocket can almost exclusively be explained by the closed and rigid structure of the helix consisting of residues 40-47 in the *Er*CRY4a protein. Most dominantly, Phe41 and Tyr107 (purple atoms in Figure 4C) would clash with 8-HDF if it was placed in the expected binding pocket. In the reference simulations of *At*CRY3 and *Xl*6-4PL, the corresponding helix loops (Figure 4C, green structure) were open.

The results of computational studies involving the MTHF ligand are summarized in Figure 5. First, the key binding residues were analyzed (Figure 5A). For the *At*CRY3 reference structure, the crystal structure revealed key contacts with residues 149, 150, 417, and 423. The binding patterns of *Deoxyribodipyrimidine* photolyase (*De*PL) and *Arthrospira platensis* photolyase (*Ar*PL) are similar, which supports the assumption that the overall fold of the binding pocket is conserved across these structures. In contrast, *Agrobacterium tumefaciens* photolyase (*Ag*PL) displayed an entirely different binding pattern for MTHF, which was dominated by backbone contacts (Fig.5A, third row). Thus, there are at least two possible and entirely different pocket shapes that display tightly packed and structurally verified binding to MTHF. The sequence of *Er*CRY4a lacks the acidic residues at the alignment positions 149 and 150 that coordinate the deeply buried pterin moiety of MTHF in the *At*CRY3 and *Ar*PL structures. Y423, which exhibits π-stacking with the benzene ring of MTHF in *At*CRY3 and *Ar*PL, is also absent in the *Er*CRY4a structure. Differences between the *Ag*PL and *Er*CRY4a sequences do not support the assumption that *Ag*PL and *Er*CRY4a share a comparable binding pattern.

**Figure 5:**
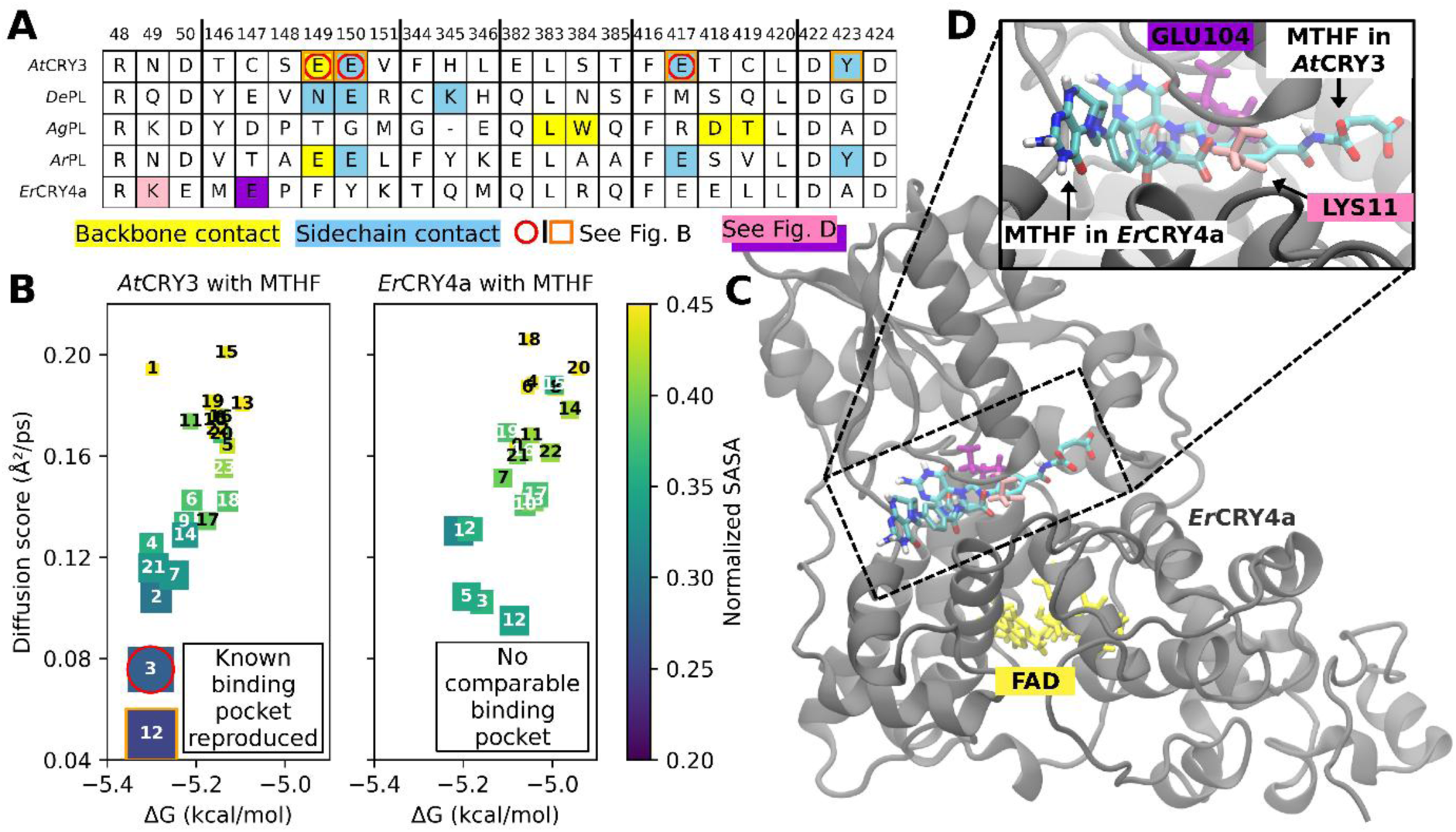
Binding properties of MTHF in the selected proteins. A: Sequence alignments of other MTHF-binders and *Er*CRY4a to *At*CRY3. The numbers in the first row enumerate the *At*CRY3 residues. B: Properties of the docking position clusters. The box sizes were scaled with the sixth power of the entropic barrier score. The text color of the cluster numbers has no significance. C: Rendering of the *Er*CRY4a conformation with MTHF docked in cluster 3. The MTHF position of cluster 3 in *At*CRY3 is shown in an overlay. D: Zoom-in of C. The sidechains of *Er*CRY4a Lys11 (Asn49 in *At*CRY3) and *Er*CRY4a Glu104 (Cys147 in *At*CRY3) are shown in pink and purple, respectively.

The results of exhaustively docking MTHF are shown in Figure 5B. For the *At*CRY3 structure, clusters 3 and 12 optimized all analyzed properties simultaneously. The docking positions of cluster 12 were in contact with all experimentally found contact residues (see orange squares in Figure 5A and B). For *Er*CRY4a on the other hand, none of the clusters show pockets with comparable properties. While the ΔG estimate of the best-performing binding pockets in *Er*CRY4a was reduced by only 0.1 kcal/mol compared to correctly identified pockets from the known binders, all other quantities were significantly worse (see Tab. 1). Via the accumulation of additional biding-related measurements, it can therefore be concluded that MTHF does not bind to *Er*CRY4a. The increased diffusion score, for example, suggests that the local contact surfaces between MTHF and *Er*CRY4a are expected to be subject to large fluctuations and faster water dynamics, which reduces the ligand retention time.

**Table 1:** Overview of the analyzed properties of the clusters with the most promising binding properties. Presence: Percentage of protein conformations in which the potential binding site (cluster) contained the docked ligand at least once after all docking attempts. ΔG: Binding score from VINA. Diffusion: Average speed of water molecules when located in the potential binding site (cluster). Normalized SASA: Ratio of molecular surface that is not covered by the protein. Entropic barrier: Measurement of blocked exit pathways out of the cluster position.

| Protein | Ligand | Cluster | Presence<br>% | $\Delta G$<br>(kcal/mol) | Diffusion<br>($\text{\AA}^2/\text{ns}$) | Normalized<br>SASA | Entropic<br>Barrier |
| --- | --- | --- | --- | --- | --- | --- | --- |
| *AtCRY3* | MTHF | 3 | 66 | -5.3 | 76 | 0.27 | 0.18 |
|  |  | 12 | 26 | -5.3 | 50 | 0.25 | 0.18 |
| *ErCRY4a* | MTHF | 3 | 70 | -5.2 | 102 | 0.36 | 0.14 |
|  |  | 12 | 26 | -5.1 | 95 | 0.34 | 0.15 |
| *Xl6-4PL* | 8-HDF | 1 | 78 | -5.6 | 50 | 0.20 | 0.19 |
|  |  | 6 | 35 | -5.3 | 38 | 0.20 | 0.19 |
| *ErCRY4a* | 8-HDF | 9 | 21 | -5.6 | 50 | 0.18 | 0.19 |
|  |  | 19 | 08 | -5.4 | 83 | 0.21 | 0.19 |

To investigate the blockage of the hypothetical MTHF binding pocket, binding positions from cluster 3 of *At*CRY3 and cluster 3 of *Er*CRY4a are shown together in Fig. 5C/D. The position of MTHF expected in *At*CRY3 served as analogous to place MTHF in *Er*CRY4a, but an ionic bond between Lys11 and either of Glu102 or Glu104 disrupts the access and positioning in the correct place. At least one of those ionic bonds was always formed and across all replica simulations. The *Er*CRY4a protein conformations were combined with docked positions of MTHF docked to *At*CRY3 and steric clashes were counted. While the ionic bond between Lys11 and Glu102/Glu104 could in principle reorient to accommodate MTHF, the helix formed by residues 40-47 and the hydrophobic residues Phe106 and Tyr107 sterically prevented the inclusion of MTHF.

## Materials and Methods

### Molecular cloning of plasmids

A pCold*Er*CRY4a plasmid was already available in the lab and it was cloned previously [15]. The full-length coding sequence for *Xl*6-4PL GenBank: AB042255.1) was synthesized by Integrated DNA Technologies (IDT) with flanking sequences (5′–ACGGCGGTTCCTACTAGT–3′ at the N-terminus and 5′–CCCGAGGGATCCGAATT–3′ at the C-terminus) to enable seamless cloning. The DNA fragment was inserted into the pCold expression vector with a His tag using the In-Fusion® HD Cloning Kit (Takara, Shiga, Japan). The plasmid, designated pCDFDuet-His_6_*FbIC* was generously provided by Prof. Dr. Lars Oliver Essen (Philipps-Universität Marburg). This plasmid contained the *Streptomyces coelicolor* (UniProt entry Q9KZZ7) *SCO4429* (*fbiC*) gene that enable the heterologous expression of the 8-HDF synthase in *E. coli*. This approach allowed us to produce this enzyme in a well-characterized and easily manipulatable bacterial host.

### Expression and purification of *Er*CRY4a and *Xl*6-4PL

Competent *E. coli* BL21(DE3) cells were transformed with pCold*Er*CRY4a or pCold*Xl*6-4PL and the transformants were selected on LB plates containing ampicillin (100 µg mL^−1^). A single colony was inoculated into 30 mL of LB media supplemented with ampicillin (100 µg mL^−1^). The overnight culture, incubated at 37 °C, was subsequently inoculated into 700 mL LB medium containing ampicillin (100 µg mL^−1^). The culture was incubated at 37 °C, 200rpm with agitation until an OD600 of 0.6 was reached. IPTG was added to a final concentration of 10 µM, and the culture was incubated at 15 °C, 160rpm for 44 h.

Antenna chromophores that bind during expression remain tightly associated with the protein throughout affinity purification[33]. So, for 8-HDF binding study, *Er*CRY4a and *Xl*6-4PL that is co-expressed with 8-HDF synthase were purified using Ni-NTA affinity chromatography only, as this purification level is sufficient for evaluating chromophore incorporation during co-expression So, for a better comparison, *Er*CRY4a and *Xl*6-4PL expressed without 8-HDF synthase were also purified using Ni-NTA affinity chromatography only. For the MTHF binding experiments, *Er*CRY4a was purified using a two-step purification workflow consisting of Ni-NTA affinity chromatography followed by anion exchange chromatography based on the surface charge of the proteins using the ÄKTA pure 25 M system (GE Healthcare, Sweden). Higher protein purity is necessary for *in vitro* reconstitution assays and for downstream biophysical characterization. All purification procedures followed previously established protocols described in Xu et al. (2021) except that 50mM Imidazole was used instead of 20mM, in the wash buffer[15].

### Co-Expression of 8-HDF synthase with *Er*CRY4a and *Xl*6-4PL

For co-expression of *Er*CRY4a and *Xl*6-4PL together with the 8-HDF synthase the same expression protocol was used with following modifications: *E. coli* BL21(DE3) competent cells were transformed with pCDFDuet-His_6_*FbIC* and pCold*Er*CRY4a or pCold*Xl*6-4PL, and streptomycin (50 µg mL^−1^) was added in addition to ampicillin in every step, due to the streptomycin resistance present on pCDFDuet-His_6_FbIC.

*Er*CRY4a and *Xl*6-4PL proteins co-expressed with 8-HDF synthase were purified by Ni-NTA affinity chromatography according to the protocol established by Xu et al. (2021) except that 50mM Imidazole was used instead of 20mM, in the wash buffer [15].

### *In vitro* reconstitution of *Er*CRY4a with MTHF

To reconstitute MTHF with *Er*CRY4a protein, 800 μM MTHF (Cayman Chemicals) dissolved in 20 mM sodium phosphate buffer (pH 7.5) was added to 16μM *Er*CRY4a. The mixture was incubated for 1 h in the dark at 4°C. The solution was then washed twice using a 30 kDa centrifuge filter and spun at 4000 x g for 7 min at 4 °C. The samples were analyzed by UV-Vis spectroscopy before and after washing using an Agilent Cary 60 UV-Vis spectrophotometer (Agilent Technologies, USA), covering a wavelength range of 200 nm–800 nm.

### Isothermal titration calorimetry (ITC) assay

ITC assay was employed to assess the potential binding interactions between *Er*CRY4a and MTHF using a VP-ITC from MicroCal (Northhampton, MA). A detailed step-by-step operation procedure of an ITC experiment was described previously [34]. For the present application we used an ITC buffer containing 20 mM sodium phosphate at pH 7.5 for preparation of stock solutions of purified *Er*CRY4a and MTHF (Cayman Chemicals). Prior to titration, all solutions were filtered and degassed using a vacuum pump, and both *Er*CRY4a and MTHF were placed in the 20 mM sodium phosphate buffer at pH 7.5 to mitigate potential artifacts arising from buffer mismatch. In each experiment, 1.6 mL of *Er*CRY4a was placed in the calorimetric cell at a final concentration ranging from 3 to 30 µM, depending on the titration. The syringe contained 300 µL of MTHF at a concentration of 400-800 µM. The titration involved 26 injections, beginning with an initial 2 µL injection, followed by 25 injections of 10 µL each. The reference power was set at 10 µcal/sec, the stirrer rate was 300 rpm, and the spacing between the injections was adjusted to allow for complete heat return to the baseline. The experiments were conducted at both 12°C and 25°C. Control titrations of MTHF into the buffer were conducted under the same conditions to account for dilution and background heat.

### MD simulations

In this study, three protein structures were subjected to MD simulations and molecular docking with the target ligands, MTHF and 8-HDF. As a positive control, MTHF-binding *At*CRY3 and 8-HDF-binding *Xl*6-4PL were used. *At*CRY3 has previously been crystallized in the presence of MTHF [22]. This crystal structure was used for simulations. The other protein structures were obtained as unrelaxed predictions of the OpenFold [35] implementation of the AlphaFold2 [36] algorithm. Structure predictions relied on the sequences for *Er*CRY4a and *Xl*6-4PL. Accurate structure predictions were possible because OpenFold had access to template structures available through the following PDB IDs: 6FN0 [animal-like Cryptochrome] [37], 1QNF [bacterial photolyase][38], 1TEZ [bacterial photolyase][39], 4U63 [bacterial photolyase] [40], 6PU0 [pigeon cryptochrome 4] [14], 6PTZ [pigeon cryptochrome 4] [14], 5ZM0 [animal-like cryptochrome] [37], 2WQ7 [drosophila 6-4 photolyase][41], and 3CVV [drosophila 6-4 photolyase] [41].

The MTHF binding pocket of the *At*CRY3 crystal structure does not involve experimentally unresolved terminal residues. Thus, it was assumed that the exclusion of flexible terminal residues did not change the relevant part of the molecular binding calculations. Additionally, the conformational space of larger flexible parts would not be explored appropriately during the planned simulation time. Excluding the flexible termini of the protein from the simulation therefore avoids artifacts that may arise due to the arbitrary initial position of the involved atoms. The resolved amino acids of the *At*CRY3 crystal structure were used as a *priori* benchmark for sufficient rigidity. After sequence-structure alignment using STAMP[42], interfaced through the MultiSeq tool [43] in VMD [44], residues that aligned with the resolved part of *At*CRY3 were retained. These included residues 39-496 for *At*CRY3, residues 1-474 for *Er*CRY4a and residues 1-475 for *Xl*6-4PL. All structures were solvated in a 120 Å cubic simulation box filled with water at a salt-concentration of 0.15 mol/L NaCl. pKa values of the protein residues were estimated with PROPKA3 [45] and two separate structures were prepared according to macroscopic pH values of 7 and 8. Studying multiple protonation states allows the exploration of different protonation-dependent conformations of proteins [46] and reduces the sensitivity to errors in the protonation state prediction tool. When the results were averaged over both pH values, an experimental pH of 7.5 was modelled.

MD simulations were performed using NAMD3 [47,48] employing the CHARMM force field [49–51] and FAD parameters [52–54]. Employing the same settings as described previously [55,56], an initial 20,000 gradient descent (standard numerical optimization algorithm) steps were followed by a 10 ns equilibration simulation using the NPT statistical ensemble with a 1 fs integration time step, followed by 400 ns simulations with a 4fs time step within the NVT statistical ensemble. From the full simulation time, the first 210 ns were identified as equilibration via RMSD convergence, leaving 200 ns for the analysis and docking calculations. Three independent replica simulations were performed for both pH values.

### Molecular Docking

As the first step in ligand preparation, potential tautomer (different protonation states), were explored. Due to the inability of crystallography to capture hydrogen and the high relevance of hydrogen bonds in molecular docking, the molecular protonation pattern should be treated carefully. Possible tautomer were manually identified and optimized in vacuum at the MP2/cc-pVTZ level of theory using the Gaussian16 software [57]. The energies were then evaluated, including an implicit solvation that solves a Poisson equation using the electronic density (SMD)[58]. For MTHF, the Pteridine molecule was used as a reduced model structure because the protonation of the remaining molecular parts (benzene, carbonylamino, and carboxyl groups) was clear. For 8-HDF, deazaisoallaxine was subjected to calculations to determine the unclear protonation states of 8-HDF.

The ligands were prepared using Meeko [59] with Gasteiger charges and explicit water molecules (hydrated docking process). To prepare the receptor structures, one frame every 400 ps was extracted from the production MD simulation and prepared with Meeko while retaining the CHARMM partial charges and maintaining FAD as part of the receptor. A custom script was used to convert the atom types of FAD into Sybyl atom types. To avoid bias in the known binding pockets, exhaustive docking was performed, including the full receptor in the search space. By aligning all structures to the *At*CRY3 crystal structure before docking, it was possible to use the same search space for all receptors and conformations, which resulted in a box of 90 × 65 × 65 Å^3^. Vina [60] was used for docking with 384 docking attempts (exhaustiveness), a grid spacing of 0.375 Å, a minimum RMSD between the output positions of 2 Å, and an energy range of 10 kcal/mol. In total, resulting in

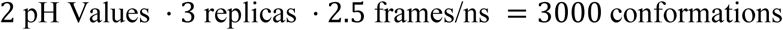

per receptor which were each subjected to 364 docking attempts. With up to 80 docked positions per docking run (some produced less), approximately 480k docked complexes were generated for each receptor-ligand pair. These structures were used to sample the conformational space of possible ligand-docking locations across the protein and were further analyzed.

### Analysis of docking positions

This section defines all the properties that were employed to analyze, rank, and characterize the binding pockets of the studied proteins.

### Alignment to known binding proteins

The crystal structures of proteins that included a bound MTHF

- *At*CRY3 [pdb ID 2IJG] [22]

- [*De*PL; 1DNP] [61]

- [*Ag*PL; 4U63] [40]

- [*Ar*PL; 6KII] [62] or 8-HDF ligand

- [*An*PL; 1QNF][38]

- [*Th*PL; 2J07] [63]

- [*Cra*CRY; 6FN2] [37]

- [*Sy*PL; 8IJY] [64]

were manually inspected, and contacts of the ligand molecules with side chains or backbones of certain amino acid residues were noted separately. Structure-informed sequence alignment using STAMP[42] was performed with the crystal structures and predicted folds of *Xl*6-4PL and *Er*CRY4a. The aligned sequences were then combined with information on key contacts.

### Clustering

For a given receptor-ligand pair, the ligand docking positions were clustered based on the most dominant contact residues. Contact residues were defined as protein (or FAD) residues that appeared within 4 Å of the ligand binding site. First, the ligand-receptor contact probabilities of all residues were computed. All docked positions where the most probable contact residue appeared to be a contact residue were assigned to cluster 1. Subsequently, the contact probabilities were recomputed using all unassigned docking positions. From the recomputed contact probabilities, all docked positions where the new most probable contact residue was a contact residue were assigned to cluster 2. This procedure was repeated until 95% of all docked ligand positions were assigned to a cluster. All remaining docked positions were assigned to the final cluster.

### Presence

The presence score for a cluster is given as the fraction of MD simulation frames in which the given cluster was found at least once within the docked positions. For visualization purposes, this property is computed every 0.1 ns but a time-average of a moving window of 5 ns is shown. Such an average is necessary since a cluster can only be either present among the docked positions for a particular simulation frame (1) or not (0), which leads to large fluctuations in the data when showing the presence over time.

**ΔG:** For the binding free energy estimate of ligands’ docked positions, the Vina score was used and termed ΔG throughout.

### Diffusion

The diffusion scores model the expected dynamics of a docked ligand based on the dynamics of the water molecules from the MD simulation. A diffusion score was assigned to each protein residue; the diffusion score of the ligands’ docked positions was computed as the average diffusion score across the contact residues. To assign the per-residue diffusion scores, the displacements over 200 ps of all water molecules within 4 Å of a residue were computed such that each water molecule within 4 Å contributed a data point. These computations were accumulated into a histogram of displacements for each residue of the protein. For a truly diffusive process, the histogram is expected to approximate a Maxwell distribution profile[65] with a single diffusion coefficient:

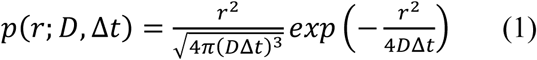

Here, the Maxwell distribution p(𝑟) is a probability distribution of the displacement *r* and depends on the time interval between observations Δt and the diffusion coefficient D. Because the dynamics of bound water molecules represent a sub-diffusive process [66,67], it is generally necessary to fit multiple Maxwell components simultaneously. For two components, the multi-component Maxwell distribution is expressed as

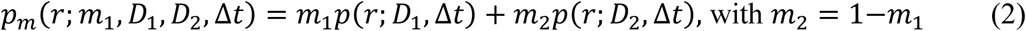

This combination of two Maxwell components allows the separation of the contributions of close but unbound water molecules and close, bound water molecules. For most residues, three components were sufficient to obtain good (R^2^ > 0.9) fit to the histograms of the mean squared water molecule displacement. For the exceptional cases, the histogram was unsteady, and the protein residues were buried, indicating a lack of data points for the histograms and not a lack of model expressiveness. Even in these cases, the fits still agreed (R^2^ > 0.8) with the overall shapes of the histograms. The diffusion score for a protein residue was computed as the average diffusion coefficient 𝐷 = ∑_𝑖_ 𝑚_𝑖_𝐷_𝑖_ .

**Normalized SASA:** For every docked position, the solvent-accessible surface area (SASA) was measured in VMD [44] using a probe radius of 1.4 Å with the surface restricted such that the contact surface between the protein and the ligand was excluded (protein+ligand as the main selection and using the ligand for the restricted option). The obtained SASA values were then normalized by the surface area obtained when the contact with the protein was ignored.

**Entropic barrier score:** The entropic barrier score is a crude estimate of the size of the exit pathway for the molecule out of the pocket. To understand the approach, one may imagine that all atoms of the ligand emit light rays, and it is measured how many of those are absorbed by the protein. Algorithmically, this is solved by selecting a random atom of the protein at position 𝑟⃗_0_ and then defining a cylinder of width R that starts at 𝑟⃗_0_ and points infinitely long into a random direction 𝑢⃗ . Subsequently, it is determined whether any atom of the protein lies within the cylinder. Repeating the calculations with many randomly selected ligand atoms and random directions allows convergence towards a score between 0 (no hits) and 1 (every cylinder contains atoms of the protein). To achieve this, one tests whether there are atoms of the protein for which the position vector 𝑟 satisfies the following conditions:

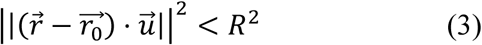

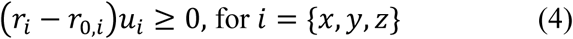

Here,”·” denotes the dot-product. Note that 𝑢⃗ is a normalized vector that is uniformly sampled from the unit sphere. The first condition ensures that the atoms are within the radius of the cylinder, and the second condition ensures that the cylinder begins at position 𝑟⃗_0_ and stretches to infinity from there. The random atoms and directions were sampled until the change in the hit rate converged below 0.01. The cylinder radius was set to R = 1.5 Å. A high entropic barrier score is expected to be strongly correlated with a high normalized SASA, but the entropic barrier score also allows differentiation between shallow binding pockets (where most light rays will miss) and deeply buried binding pockets (where most light rays will eventually hit a protein atom).

**Clash probability:** The protein folds were overall similar, but the locations of the docked ligands were different. A measurement was desired to count the number of atoms that would clash if the docked position of a chromophore inside a protein was overlayed with a different protein structure. For this purpose, the docked positions of MTHF in *At*CRY3 and 8-HDF in *Xl*6-4PL that were found within the most favorable binding clusters were selected. For each docked ligand position in the reference proteins, it was counted how often a heavy atom from the *Er*CRY4a trajectory was within 1 Å of a heavy atom of the ligand. Using the pre-aligned positions generated for docking preparation, the resulting clashes were directly attributed to differences in the internal coordinates of the proteins. The number of frames showing a clash with any atom of a certain residue was normalized by the total number of simulation frames of the *Er*CRY4a trajectory, resulting in a clash probability for every residue.

## Discussion

Cryptochrome 4a binds the chromophore FAD and is believed to play a crucial role in the ability of birds to detect the Earth’s magnetic field for navigation[3,4,11–15,55]. To date, it is unknown whether CRY4a binds to any other secondary chromophore. Here, we studied the interaction of *Er*CRY4a with two commonly reported chromophores, MTHF and 8-HDF. Co-expression of *Er*CRY4a with 8-HDF synthase did not show the characteristic absorption peak at 440nm which was otherwise present in the co-expression of the positive control protein *Xl*6-4PL with 8-HDF synthase. The absence of this absorption peak (440nm) provides compelling evidence that 8-HDF does not bind to *Er*CRY4a. Due to the low solvent accessibility of the expected binding pockets (see section 3.3), loss of 8-HDF due to the purification process is unlikely. These results render it very unlikely that 8-HDF binds to *Er*CRY4a, indicating a significant difference in the chromophore-binding properties between *Er*CRY4a and *Xl*6-4PL. For MTHF, both *in vitro* reconstitution experiments and isothermal titration calorimetry (ITC) consistently showed the absence of stable association between MTHF and *Er*CRY4a. Our experimental findings therefore strongly suggest that *Er*CRY4a does not use either of these common chromophores as antennae and suggest that FAD alone is responsible for the light-sensing and likely function of *Er*CRY4a as the primary magnetic sensor.

In our computational analysis, we found no similarity at the relevant binding sites between the sequences of *Er*CRY4a with proteins shown to bind MHTF or 8-HDF. This suggests that the binding pocket is not conserved, leaving the possibility of discovering a novel binding conformation. The approach to combine protein ensembles from MD simulations with exhaustive docking, careful preparation of protonation, and water dynamics-informed ranking, reproduced the expected binding pockets for 8-HDF in *Xl*6-4PL and MTHF in *At*CRY3. Following this validation, *Er*CRY4a was investigated as a test case. No relevant binding pocket was found for MTHF, whereas a potential novel binding pocket in *Er*CRY4a was found for 8-HDF. The absence of the expected binding pockets was rationalized through analysing potential steric clashes. A rigid helix formed by residues 40-47 in *Er*CRY4a was found to be mostly responsible for preventing MTHF binding. Furthermore, the protein dynamics suggested that aromatic residues around the entrance to the identified potential novel 8-HDF binding pocket in *Er*CRY4a effectively hindered 8-HDF from entering it.

The present findings carry important implications for the validity of laboratory-based behavioural tests investigating magnetoreception in migratory birds under the radical pair hypothesis [68–72]. European robins were captured from the wild and subsequently maintained for extended periods under indoor housing conditions prior to behavioural testing. The laboratory diet, which consists primarily of grains, industrially produced mealworms, and vitamin supplements, lacks the algae and bryophytes that synthesize 8-HDF [26–28]. Birds kept long-term under these conditions would therefore most likely be depleted of 8-HDF. However, *Er*CRY4a did not bind 8-HDF, suggesting that it is not a required cofactor for *Er*CRY4a dependent magneto sensory processes. Accordingly, a deficiency of 8-HDF in captive birds does not compromise laboratory magnetoreception studies. Regarding MTHF, which also appears to be unlikely to be bound with CRY4 protein, birds should be able to synthesise MTHF from dietary folates. The European robin genome (bEriRub2.2; GCA_903797595.2) contains genes encoding the key enzymes, including MTHFD1, MTHFD2L, SHMT1, and SHMT2. Taken together, these considerations resolve the long-standing concern regarding the nutritional availability of CRY4 secondary chromophores in behavioural experiments using captive migratory birds.

Our combined experimental and computational results suggest that neither 8-HDF nor MTHF associates with *Er*CRY4a. These findings suggest that FAD alone serves as the catalytic chromophore in *Er*CRY4a. Although our study provides strong evidence against the binding of 8-HDF and MTHF to *Er*CRY4a, it does not exclude the possibility that other, yet unidentified, secondary chromophores or cofactors could be involved. Therefore, we suggest that future studies on tissue-derived native *Er*CRY4a should be performed to confirm the absence of additional chromophores or potentially discover novel antenna chromophores.

## Author Contributions

A.B.P.A. conceived the study, designed and performed the experiments, analysed and interpreted the data, and wrote and edited the manuscript. B.D. performed ITC experiments together with A.B.P.A., contributed to data analysis, and assisted in writing the manuscript.

J.H. performed the computational analyses and contributed to writing the manuscript. T.K., and R.B. conceived parts of the study and provided scientific advice and mentoring. G.D., G.S., and J.S. assisted with protein expression and purification. J.X. provided scientific advice and mentoring throughout the project. I.A.S., K.W.K., and H.M. supervised the project and contributed to data interpretation, manuscript revision, and constructive discussions. All authors read and approved the final manuscript.

## Acknowledgements

This research was supported by the Deutsche Forschungsgemeinschaft (DFG) through the Collaborative Research Centre “Magnetoreception and Navigation in Vertebrates” (SFB 1372, grant no. 395940726 to IAS, KWK, and HM), through the Transregio Research Centre “HYP*MOL” (TRR386, grant no. 514664767 to IAS) and through the Excellence Cluster “NaviSense” (EXC 3051, grant no. 533653176 to TK, IAS, HM). The research was also supported by the European Research Council (ERC) under the European Union’s Horizon 2020 research and innovation program (Grant Agreement No. 810002, Synergy Grant “Quantum Birds”). The authors also thank the Volkswagen Foundation (Lichtenberg Professorship awarded to IAS), the Ministry of Science and Culture of Lower Saxony Simulations Meet Experiments on the Nanoscale: Opening up the Quantum World to Artificial Intelligence (SMART), and Dynamik auf der Nanoskala: Von kohärenten Elementarprozessen zur Funktionalität (DyNano). The authors gratefully acknowledge the computing time granted by the Resource Allocation Board and provided on the supercomputer Emmy/Grete at NHR-Nord@Göttingen as part of the NHR infrastructure. The calculations for this research were conducted with computing resources under the project nip00058. We thank Prof. Dr. Lars Oliver Essen (Philipps-Universität Marburg) for kindly providing the pCDFDuet-His₆-FbiC plasmid as a gift. We also thank all members of the Mouritsen group for their support and discussions during this work.

## Data availability

The data referred to in this manuscript are present in the manuscript and in the supporting information. Unprocessed data are deposited on a server of the University of Oldenburg in accordance with the data policy of the Collaborative Research Centre SFB 1372 and can be available on request from the corresponding author.

## Abbreviations

The abbreviations used are: CRY4a,Cryptochrome 4a; *Er*CRY4a, *Erithacus rubecula* cryptochrome4a; 8-HDF, 8-hydroxy-5-deazaflavin; MTHF, 5,10-methenyltetrahydrofolate; *Xl*6-4PL*, Xenopus laevis* 6-4 photolyase; ITC (Isothermal Titration Calorimetry); MD, Molecular Dynamics; SASA, Solvent Accessible Surface Area; *At*CRY3, *Arabidopsis thaliana* Cryptochrome 3; *De*PL, *Deoxyribodipyrimidine* photolyase; *Ag*PL, *Agrobacterium tumefaciens* photolyase; *Ar*PL, *Arthrospira platensis* photolyase; *An*PL, *Anacystis nidulans* photolyase; *Th*PL, *Thermus thermophilus* photolyase; *Cra*CRY, *Chlamydomonas reinhardtii* Cryptochrome; *Sy*PL, *Synechococcus elongatus* 6-4 photolyase

**Conflict of interest:** The authors declare no conflicts of interest.

