## supplemental file for "Absence of 8-HDF and MTHF Antenna Chromophore Binding in *Er*CRY4a Suggests a Possible Flavin-Only Cofactor State: Insights from Biochemical and Computational Analyses"

* To whom correspondence should be addressed.

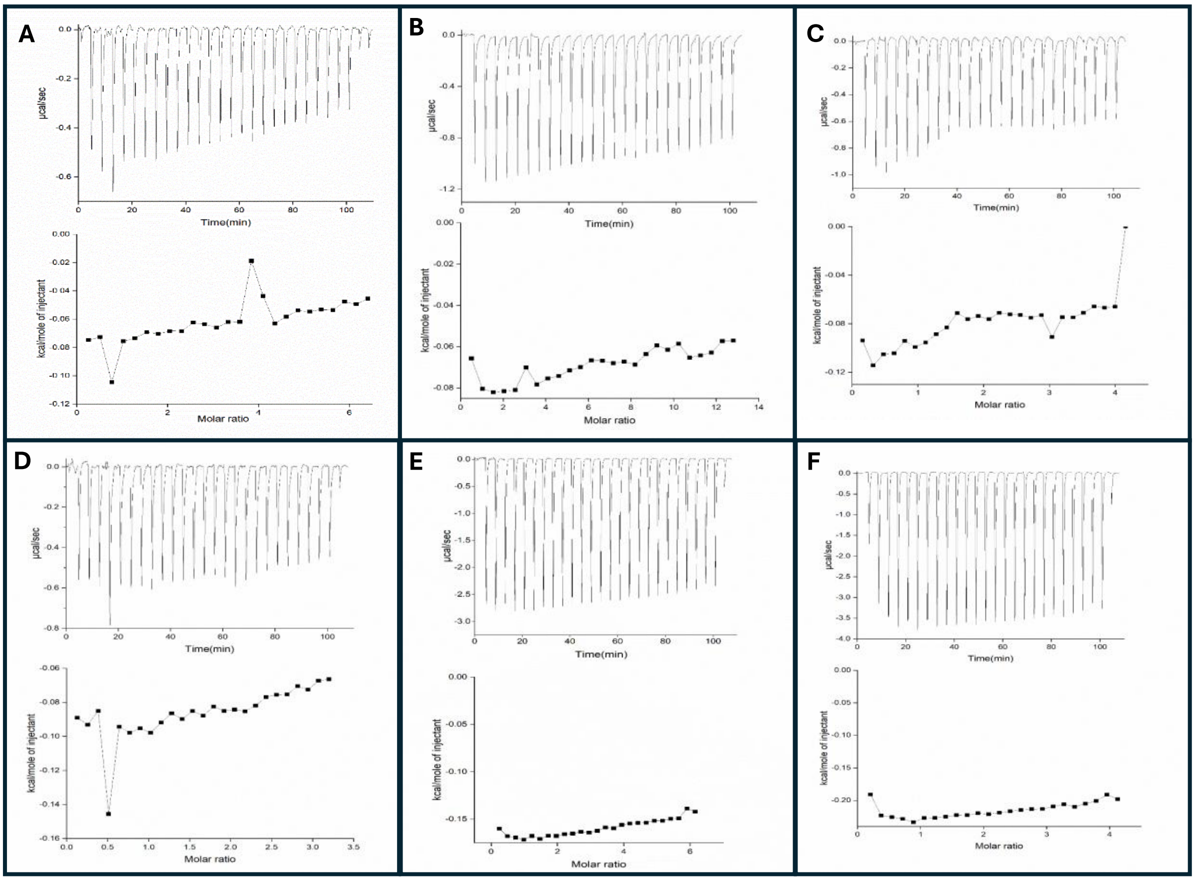


**Figure S1: ITC experiments probing *Er*CRY4a–MTHF interaction at 12 °C across protein and ligand concentrations.**

Integrated heats from ITC titrations performed at 12 °C using ErCRY4a concentrations of 10–30 µM in the cell and MTHF concentrations of 400 or 800 µM in the syringe. Panels are ordered by increasing ErCRY4a concentration and, within each protein concentration, by increasing MTHF concentration: (A) 10 µM *Er*CRY4a / 400 µM MTHF, (B) 10 µM / 800 µM, (C) 16 µM / 400 µM, (D) 20 µM / 400 µM, (E) 20 µM / 800 µM, (F) 30 µM / 800 µM. None of the tested conditions exhibited saturation or sigmoidal curvature indicative of binding. The measured heats were dominated by ligand dilution and closely resembled ligand-to-buffer control experiments; therefore, data are shown without subtraction to avoid artifacts arising from dilution heats exceeding the total signal.

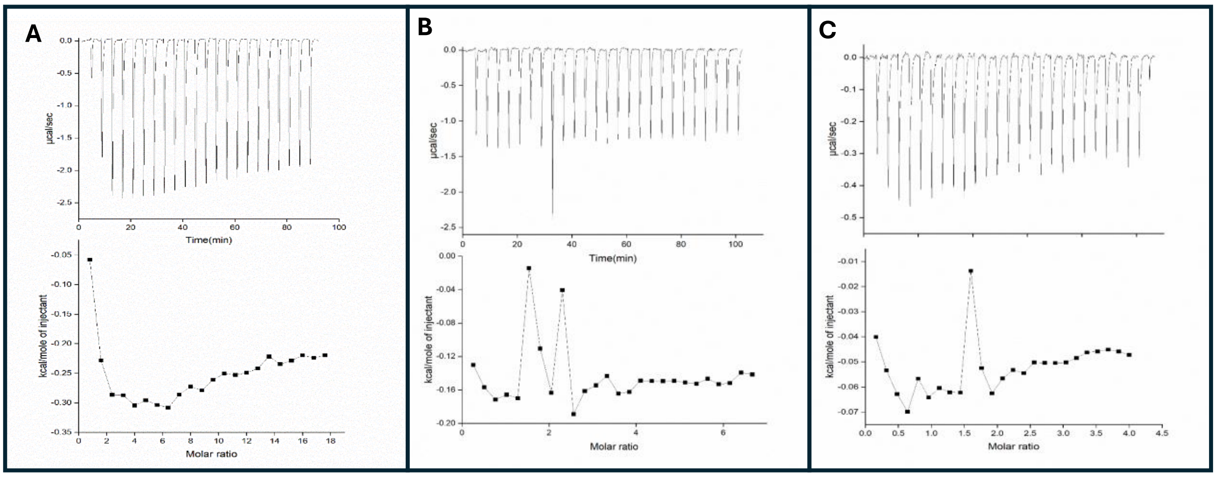


**Figure S2: ITC experiments probing *Er*CRY4a–MTHF interaction at 25 °C across protein concentrations.**

Integrated heats from ITC titrations performed at 25 °C using *Er*CRY4a concentrations of 3–16 µM in the cell with a constant MTHF concentration of 400 µM in the syringe. Panels are ordered by increasing protein concentration: (A) 3 µM *Er*CRY4a / 400 µM MTHF, (B) 10 µM / 400 µM, (C) 16 µM / 400 µM. None of the tested conditions displayed saturation behavior or sigmoidal curvature indicative of binding. As observed at 12 °C, the measured heats were dominated by ligand dilution and were comparable to ligand-to-buffer controls; data are therefore presented without subtraction.

**Table S1. Summary of ITC experiments performed to probe *Er*CRY4a–MTHF interaction across temperatures and concentrations**

ITC measurements were conducted at 12 °C and 25 °C over a range of *Er*CRY4a (3–30 µM) and MTHF (400–800 µM) concentrations to maximize the likelihood of detecting a binding isotherm.

| **Temperature** | **[*Er*CRY4a] (µM)** | **[MTHF] (µM)** | **Outcome** |
|---|---|---|---|
| 12 °C | 10 | 400 | no saturation |
| 12 °C | 10 | 800 | no saturation |
| 12 °C | 16 | 400 | no saturation |
| 12 °C | 20 | 400 | no saturation |
| 12 °C | 20 | 800 | no saturation |
| 12 °C | 30 | 800 | no saturation |
| 25 °C | 3 | 400 | no saturation |
| 25 °C | 10 | 400 | no saturation |
| 25 °C | 16 | 400 | no saturation |
